# Sexual Dimorphism of Cancer-Associated Fibroblasts Governs Matrix and Vascular Organisation in Breast Cancer

**DOI:** 10.64898/2026.08.27.747484

**Authors:** Peng Liu, Fiona R. Saunders, Matthew Everest, Natthaya Eiamampai, Matthew P. Humphries, Camilla Coulson Gilmer, Giulia Conti, Lucy F. Stead, Rasha Abu-Eid, Valerie Speirs

**Affiliations:** School of Medicine, Medical Sciences and Nutrition, University of Aberdeen, Aberdeen AB24 2ZD; National Pathology Imaging Cooperative, Leeds Teaching Hospitals NHS Trust, Leeds LS9 7TF; Manchester Cancer Research Centre, University of Manchester, Manchester, M20 4GJ; Department of Experimental and Clinical Medicine, Azienda Ospedaliero-Universitaria Careggi, University of Florence, Florence, Italy; Leeds Institute of Medical Research at St James’s, University of Leeds, Leeds LS9 7TF; School of Dentistry, School of Health Sciences, College of Medicine and Health, University of Birmingham, Birmingham B5 7EG; Aberdeen Cancer Centre, University of Aberdeen, Aberdeen AB25 2ZD

**Keywords:** breast cancer, sex, CAFs, transcriptomics, tumour microenvironment

## Abstract

Breast cancer (BC) shows greatest sexual diversity. Increased diagnosis and poorer outcomes in men highlights the need to better understand its biology. We hypothesised that cancer-associated fibroblasts (CAFs), the most abundant cell type in the tumour microenvironment, might define sex-related differences.

Using phenotypically matched male and female CAFs generated from breast cancer tissues, we demonstrate distinct transcriptional programmes and functional behaviours associated with extracellular matrix remodelling, cell adhesion, migration and vascular development. Compared to CAFs generated from females BC, those from males generated denser, more complex matrices promoting stronger tumour and endothelial cell adhesion, vascular growth, but less organised capillary network formation.

Findings reveal fundamental sex-related variations in CAF phenotype and biology in BC. These findings highlight the need to integrate biological sex into precision oncology to identify opportunities for sex-specific therapeutic strategies in BC.

## Introduction

Sexual diversity is now recognised in cancer with the greatest diversity seen in breast cancer (BC) [1]. Transcriptomic [2–5], bioinformatic [6] and deep learning studies [7] have demonstrated that male and female BC are not the same, however this has been difficult to study at a functional level. Unlike the wide range of *in vitro* [8] and animal models [9] available to study BC in women, male BC models do not exist. The rare presentation of male BC, accounting for <1% of all BC cases [10], reduces availability of samples required to develop *in vitro* models, adding to this challenge. Given the rising numbers of men receiving a BC diagnosis, inferior outcomes related to late diagnosis [11], and data from the Surveillance, Epidemiology and End Results (SEER) database (2010-2022) showing that neoadjuvant therapy was less effective in men than in women with early-stage breast cancer [12], *in vitro* models are necessary to help identify potential sex-specific therapeutic targets.

The tumour microenvironment (TME) comprises a rich diversity of cell types, including cancer associated fibroblasts (CAFs) and immune cells which are embedded within a collagen-rich extracellular matrix (ECM), collectively known as the stroma [13]. Previous work has shown dissimilarity between the TME in male and female BC. For example, the manifestation of elastosis is initiated by secretion of the elastin precursor, tropoelastin, by CAFs. Elastosis did not influence outcome in male BC, but did in female BC [14]. Similarly, CD8, a marker of good outcome in female BC had the opposite effect in male BC [15] and reduced frequency of the immune checkpoint inhibitor PD-1, found on T-cells residing within the TME, has been reported in male compared to female BC [16]. Transcriptomic analysis of the immune landscape in BC also supports sex-specific immune profiles [17, 18].

Cancer cells have a symbiotic relationship with CAFs, the main cell type within the complex multicellular TME. CAFs interact extensively with cancer cells, either directly through paracrine signalling [19], extracellular vesicles [20] or by mediating cell‒cell adhesion [21], or indirectly through ECM remodelling and immune cell infiltration [22]. They also provide three-dimensional support through deposition of ECM components, additionally supporting the metabolic and nutritional requirements of cancer cells [13].

CAFs are abundant in BC with increasing evidence for their role in influencing tumour cell behaviour. For example, the tumour:stroma ratio can influence both outcome and therapeutic response in BC while CAFs themselves can influence the effects of chemotherapy [23–25]. Spatial technologies have enabled CAF subtypes to be identified across various pathologies, including breast [26, 27] creating the possibility for biomarker-driven drug development targeting CAFs.

Recognising the importance of CAFs in breast carcinogenesis, we have taken a lateral approach towards studying male BC *in vitro*. Here, we have generated and characterised CAFs from male and female BC and demonstrate how biological and functional properties related to adhesion and vascularisation are influenced by sex.

## Materials and Methods

### CAF generation

Following ethical approval (Leeds (East) Research Ethics Committee (REC);15/YH/0025) CAFs were derived directly from matched male (n=7) and female (n=6) breast tumours (ductal NST, grade 2, ER+) obtained from surgical resections or obtained as frozen stocks from the Breast Cancer Now Biobank (East of England – Cambridge Central REC; 23/EE/0229) and characterised according to a previous protocol [28]. As non-immortalised cells, these have a finite lifespan in culture, entering senescence after 5-6 passages, limiting their long-term use. Since men present infrequently for breast cancer surgery, it was necessary to build CAF stocks hence these were hTERT-immortalised [29].

For consistency, this was applied to both male and female CAFs, with one exception. CAF use across specific experiments is outlined in **STable 1**, with at least 2 different CAFs per sex used in each experiment. With no single biomarker that defines CAFs unconditionally [30], we isolated CAFs directly from surgically resected BCs and characterised them based on spindle shaped morphology and used a panel of positive and negative biomarkers defined in the literature to identify these cells by morphology, immunofluorescence and gene expression (29, 43). This included negativity for epithelial (EMA) and endothelial (CD31) markers, and positivity for mesenchymal markers vimentin and α-SMA, using immunofluorescence, as described previously [31]. Following RNA-seq analysis (described below), CAFs were further categorised based on published gene expression profiles for CAFs, covering the major cell types found in the breast cancer microenvironment [32].

### Cell culture

Cells used and their culture conditions are shown in **STable2**. All cultures were maintained at 37 °C, 5% (v/v) CO_2_. HUVECs were used at passages 4 - 9 and were routinely grown in flasks coated with 10 µg/ml gelatin (Sigma-Aldrich). Quarterly mycoplasma checks were consistently negative and annual STR profiling of non-primary cells confirmed fidelity (both Eurofins Scientific, UK). Conditioned medium (CM) was generated from CAFs. Cultures around 80% confluent were rinsed twice in PBS and incubated for 48 h with serum-free media. Media were removed, filter sterilised (0.2 μm, Minisart) and either used immediately or stored at −80°C until required. Serum-free medium, cultured under identical conditions but in the absence of cells, was used as a negative control.

### RNA-seq

Immortalised matched male and female BC CAFs were selected (n=3 per sex). RNA quality and quantity in each sample was determined using a BioAnalyser (Agilent) before 100 ng of total RNA was used to make polyA selected Illumina compatible sequencing libraries using the TruSeq stranded RNA protocol (Illumina). The libraries were quantified by qPCR, then pooled and sequenced on a single high output NextSeq 500 (Illumina) lane to generate 76 bp single end read data. The data were base-called and de-multiplexed using the bcl2fastq software (Illumina). Data generated was stratified for sex-related differences in terms of putative CAF, fibroblast, epithelial and vascular markers. As various CAF subsets have been reported CAFs were additionally stratified according to subtypes defined for those determined from female BC [33]. Genes with an adjusted *p*-value < 0.05 and an absolute log₂ fold change > 1 were considered significantly differentially expressed in male and female BC-derived CAFs; 74 genes met these criteria.

### Analysis of Gene Ontology (GO) pathways

Functional enrichment of 43 differentially expressed genes identified from the RNA-seq data was performed using GO pathway analysis in Enrichr [34]. The top 10 significantly enriched GO terms (Biological Processes 2021) and ranked by p-value were visualised in bar graphs to highlight biological processes, molecular functions and cellular components differing between male and female BC-derived CAFs.

### Organotypic angiogenesis assay

HUVEC were plated onto confluent monolayers of male or female CAFs (described above) growing in 24 well plates, according to a published protocol [35] and cultured for 14 days. Tubular vessel-like structures were visualised by immunohistochemical staining of CD31 (Cellworks ZHA-1225) according to the manufacturer’s protocol. In brief, multilayer co-cultures were fixed with ice-cold ethanol (30 min at room temperature) and labelled with a monoclonal mouse anti-human CD31 antibody (clone: JC70A, dilution 1:400) in blocking buffer (PBS plus 1% BSA) overnight at 4°C. Secondary antibody (alkaline phosphatase (AP)-conjugated goat anti-mouse;1:500 in blocking buffer) was added and incubated for 2 hours at 37°C. After rinsing in PBS and deionised water, co-cultures were incubated with chromogen (BCIP/NBT in deionized water) for 20 mins. At least 5 images per well were taken at x4 magnification (EVOS). ImageJ was used to quantify CD31-positive areas as a measure of the formation of tubular structures.

### Wound closure and motility of HUVEC in response to CAF CM

HUVECs were plated to at least 90% confluence in each chamber of a two-chamber insert on a glass-bottomed 35 mm dish (both Ibidi, Germany) and allowed to attach overnight. The gap between the chambers was 500 ± 100 µm. After removal of the chamber, 2 ml of either fresh HUVEC medium or CAF CM was added with the Hololid replacing the standard plate lid. Progressive changes in wound healing, motility and migration were analysed using label-free time-lapse digital holography (Holomonitor M4, PHI, Sweden) in a humidified CO_2_ incubator (37°C, 5% CO_2_). Images were captured at 15 min intervals over 24 h. Quantitative measurements were analysed and extracted using Holomonitor App Suite software (version 4.0), exported to Excel and analysed using GraphPad Prism (version 10.6.0). Two-sample t-tests were applied to evaluate significant differences between CAFs.

### CAF-derived matrix (CDM) production and imaging

CDMs were generated from male and female CAFs and human dermal fibroblasts HDF377 (TCS CellWorks) derived from a female donor, as described [36]. Briefly 13 mm diameter coverslips (Epredia, UK) were coated with 1% (v/v) gelatin solution (Merck, UK), cross-linked with 1% (v/v) glutaraldehyde (Merck, UK) and quenched using 1 M glycine (Merck, UK). Cells (50,000 cells/well) were seeded onto gelatin-coated coverslips in complete growth medium supplied with 10% FBS. At confluence, they were treated on alternate day with 50 µg/ml ascorbic acid (Merck, UK) in complete growth medium for 10 days before the matrix was extracted. CDMs were fixed in 10% neutral-buffered formalin for 15 minutes at room temperature before washing with PBS (Sigma, UK). Following blocking (30% horse serum in PBS, 5 minutes at room temperature) they were incubated with Collagen type I primary antibody (AB138492, Abcam, UK; 1:100 dilution in 30% horse serum) for 1 h at room temperature. CDMs were then washed (PBS, 3 x 2 min) before application of Alexa Fluor 594 secondary antibody (Fisher Scientific, UK; 1:500 30%(v/v) horse serum) for 1 h at room temperature. After incubation the CDMs were washed with PBS then a final wash with distilled water. CDMs were mounted onto slides using Mowiol mounting solution and allowed to cure overnight, protected from light. Slides were stored in the dark at 4°C prior to confocal imaging (Zeiss LSM 880 Airyscan, Zeiss, Germany).

### CDM thickness

Measurements were performed in Zen v3.11 software (Zeiss). A Z-stack of the CDM was taken using a LSM 880 Confocal Microscope (Zeiss; 40x). The saved Z-stack was viewed in Zen, where the channel display was altered to increase the pixel brightness of the image. The start and end point of the Z-stack were defined and the 3-D distance measured in µm. For each CDM, 3 separate Z-stacks were taken to ensure that the measurements were representative of the whole CDM and the mean thickness ± SD was calculated. Directionality of collagen fibres was analysed using the ImageJ plugin, TWOMBLI ([37]

### Adhesion assay

MCF-7, HB2, male MEC and HUVEC cells were serum starved overnight before plating onto the CDMs described above. CDMs were blocked for 1 h (0.5 % BSA in DMEM) at 37°C prior to chilling on ice for 1 h. Cells were seeded at 1x10^5^ cells/ml in complete growth medium for the respective cell lines. Cells plated directly onto the tissue culture plate served as controls. After adhesion (1 h, 37°C) the plates were shaken (15 seconds) to remove non-adherent cells then washed (0.1% BSA in DMEM) three times before fixing with 10% neutral-buffered formalin (15 minutes). After a further wash, plates were stained with crystal violet (5 mg/ml in 2% (v/v) ethanol, 10 minutes). Plates were then washed with distilled water, allowed to air dry for 45 minutes before lysis with 10% glacial acetic acid (v/v in dH_2_O) for 15 minutes at room temperature. 100 µl aliquots were transferred to a 96 well plate (min. 5 technical replicates per treatment). Plates were read at 560 and 700 nm (Pherastar plate-reader, BMG LabTech).

### Fractal and lacunarity analysis to define shape complexity between male and female BC

Four µ FFPE sections of matched ER-positive male and female BC (ductal NST) were dewaxed and stained with picrosirius red to detect collagen fibres as described [38] and digital images (20x) created using a digital scanner (Aperio Scan Scope XT, Milton Keynes, UK). These were subjected to colour deconvolution using QuPath [39]. To separate different stains, the staining vectors in the images were cropped. Images were then converted to binary images using Otsu thresholding in a custom batch processing script in ImageJ [40]. Box counting was used to calculate the fractal dimension of the collagen fibres in in the images. The ImageJ built in box counting plugin was used using 400x400µm boxes with resultant output saved as a spreadsheet for further analysis. To assess distribution of space between the collagen fibres, lacunarity was measured in the same binary images using the FracLac extension for ImageJ [41] to perform sliding box lacunarity analysis using the default settings.

### 3D microfluidic tube formation assay

HUVECs and CAFs (6 million cells/mL, 1:1) were seeded in a fibrin gel (2.5 mg/ml human fibrinogen; F3879, Merck) dissolved in serum-free basal Endothelial Cell Growth Medium 2. Thrombin (5 U per 10 mg fibrinogen;605206, Sigma-Aldrich) was then added to convert the soluble fibrinogen into insoluble fibrin strands. Immediately after gentle mixing, the fibrin solution was pipetted into the central chamber of a microfluidic chip (AIM Biotech 3D cell culture; DAX01, Merck) where it polymerised into a gel. EGM-2 media supplemented with enQuireBio™ Recombinant Human VEGF-165 Isoform Protein (40 ng/ml;15965689, Fisher) was then added to the two lateral flow channels. Gravity-driven flow was introduced by adding 50 µL of medium into two connected ports of the same channel and 70 µL into the opposite two connected ports. This differential volume created a transient flow across the porous fibrin gel and the anastomotic microvessels in the chip after 4 days upon daily medium changes. CD31 immunofluorescent staining was performed to visualize the formed vascular structure. Briefly, after washing the microfluidic chips with PBS, the cells were fixed with 4% paraformaldehyde for 15 minutes, permeabilized (0.1% Triton™ X-100, 10 minutes) and blocked (2% BSA, 45 minutes at room temperature). The cells were labelled with anti-CD31 Monoclonal Antibody (Gi18, eBioscience™;Product # BMS137) at 5 µg/mL in 0.1% BSA, overnight at 4°C and then labelled with Donkey anti-Mouse IgG (H+L) Highly Cross-Adsorbed Secondary Antibody (Alexa Fluor Plus 488;Product # A32766,1:400 dilution) for 45 minutes at room temperature. Cells were then counterstained with DAPI (final concentration 1μg/ml; 10236276001, Roche) and imaged (EVOS M5000 Imaging System).

### Statistical analysis

Statistical analyses were performed using GraphPad Prism (version 10.6.0; GraphPad Software, San Diego, CA, USA). Data are presented as mean ± standard deviation (SD) unless otherwise stated. Differences between groups were assessed using appropriate parametric or non-parametric tests as indicated. A p-value <0.05 was considered statistically significant.

## Results

### CAF characterisation

The pipeline for the generation and characterisation of CAFs is shown in **SFigure 1**. CAFs displayed a typical spindle-shaped morphology and were negative for epithelial (EpCAM, keratins) and endothelial (CD31, VEGF) biomarkers and positive for αSMA, vimentin and S100A4. In general, well-established CAF/fibroblast markers identified by RNA-seq showed no sex-related differences. This was reflected in qPCR validation, consistent with immunohistochemical studies of ACTA2/α-SMA expression in CAFs from male and female BC (44, 45). RNA-seq showed that the absolute log₂ fold change between male and female BC-derived CAFs for recognised epithelial and vascular markers was < 1 (**SFigure 2**). However, *DES*, a muscle-specific protein, which can be expressed in CAFs transitioning towards a myofibroblastic or smooth muscle-like phenotype (46), showed >1 absolute log₂ fold change in female compared to male CAFs, potentially reflecting phenotypic plasticity in females. When stratified for CAF subtypes, many of the genes associated with the inflammatory CAF (iCAF) phenotype were preferentially expressed in male BC-derived CAFs, except for *DES*. Reticular subtypes (rCAF) also appeared to display sex-specificity in favour of males (**SFigure 3)**.

### RNAseq and Gene Ontology (GO) analysis

To assess differential gene expression between male and female BC-derived CAFs, a volcano plot was generated (**SFigure 4a**). Genes located towards the upper left of the plot represent those significantly upregulated in male BC-derived CAFs while those on the upper right correspond to those upregulated in CAFs from female BC. Since sex-specific transcriptional profiles were observed, GO enrichment analysis performed for biological process using Enrichr [34] revealed that the top enriched terms associated with male BC-derived CAFs were primarily associated with extracellular matrix organisation/adhesion and vascular development (**SFigure 4b**). These data suggested that the differentially expressed genes in male BC-derived CAFs may be involved in cell–cell or cell–matrix interactions essential for tissue organisation and in the formation and remodelling of blood vessels. This was then explored experimentally.

### Phenotype and adhesive properties of CDMs derived from male and female breast cancer CAFs

CDMs from female BC-derived CAFs (**Fig 1a**) and cell-free matrices from female HDF377s (**Fig 1b**) displayed a looser structure than those from male CAFs (**Fig 1c**), which were more densely packed and around 30% thicker (**Figure 1d**). Coherency analysis showed greater alignment of CDMs derived from male BC-derived CAFs (**Fig 1e**). The ability of CDMs in promoting adhesion of different cell types was compared to that of standard tissue culture plastic (**Fig 1f-i**). Adhesion of female HB2 cells to all CDM phenotypes was not significantly different to that of tissue culture plastic (**Fig 1f**). Greatest adhesion was seen with male MECs, especially to male CAF-derived matrix (**Fig 1g**). MCF-7 adhesion to male CAF-derived CDMs, was significantly greater than to female CAF-derived CDMs or that of HDF377 (**Fig 1h**) while HUVECs showed greater adhesion to male CAF-derived CDMs only, compared to plastic control (**Fig 1i**).

**Figure 1:**
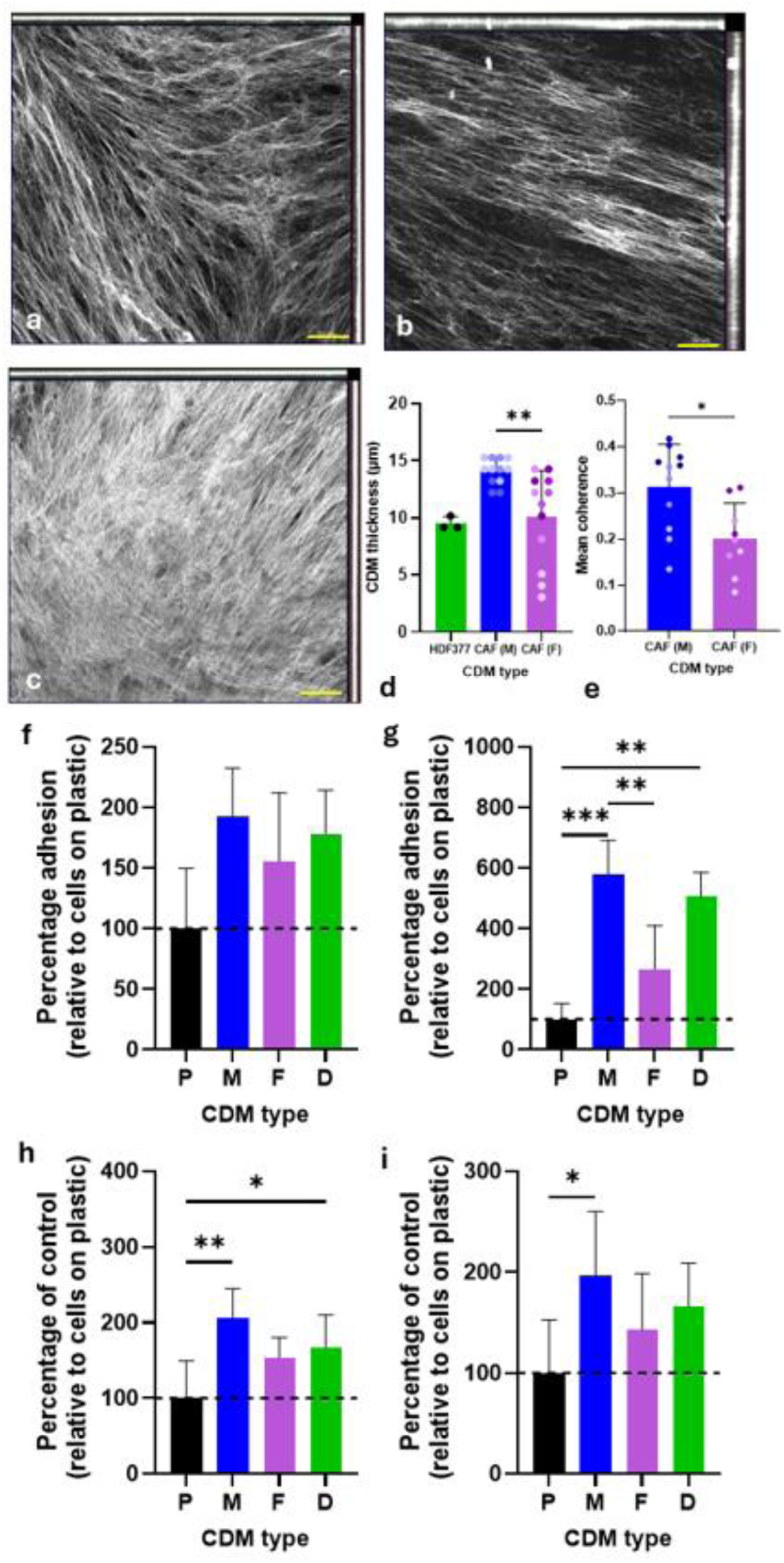
Phenotype and thickness of CDMs. CDMs derived from female CAFs (a) and HDF377 (b) had a looser structure compared to the densely packed appearance of those from male CAF-derived CDMs (c). Images are maximum intensity projections of collagen type I generated from Z-stacks. CDMs from male CAFs were also significantly thicker than those from female CAFs and HDF377 (d) and showed greater coherence (e). Adhesion of HB2 (f), MMEC (g), MCF-7 (h) and HUVEC (i) was dependent on CDM source, with MMEC showing greatest adhesion relative to tissue culture plastic. Values for individual patient samples are indicated as coloured dots. P = plastic, M = male CAF, F = female CAF, D= dermal fibroblast (377HDF). Scale bar = 50µm. *P < 0.05, **P < 0.001, ***p <0.0001

### The complexity of collagen fibres in the TME

Having demonstrated sex-dependent features in the structural and adhesive properties of CAF CDMs *in vitro*, we then explored if this translated to patient BC tissues. We used BC tissue sections stained with picrosirius red to examine the complexity of different stromal areas of the tumour microenvironment in which CAFs reside. No differences were observed between sexes in the peritumour areas (**Fig 2b, d**), but male BC tissues displayed significantly greater complexity in intra-tumoral areas. This was reflected in both fractal analysis, a measure of complexity and space filling properties and lacunarity, a measure of the heterogeneity and distribution of empty spaces within an area (**Fig 2c**).

**Figure 2:**
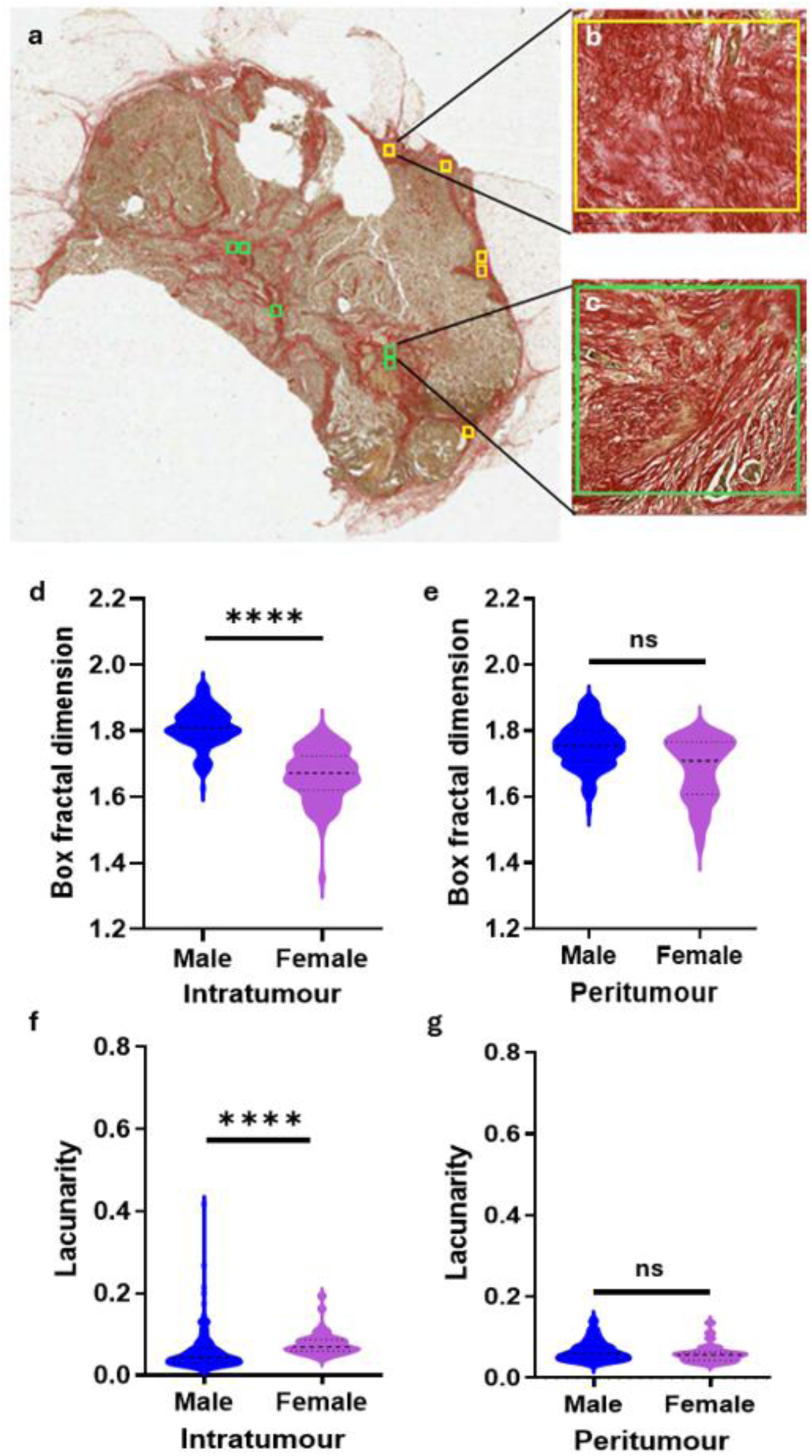
Fractal dimension analysis of peri- and intra-tumour collagen in Picrosirius red stained histological sections of ER-positive male and female breast cancer. Male BC tissue section stained with picrosirius red (a) with peri- and intra-tumour areas identified by the yellow and green boxes, respectively, expanded in b and c. No differences were observed between sexes in the peritumour areas, but in intra-tumour areas male breast cancer showed significantly greater complexity compared to female breast cancer. This was reflected by fractal analysis (a,b) and lacunarity (c,d). Bold lines on violin plots represent median/mean fainter lines show upper/lower quartiles. **** p <0.0001

### Crosstalk between CAFs and vascular cells

As vascular development was identified as another biological process strongly associated with male BC-derived CAFs we reasoned that this could indicate enhanced vascular formation and remodelling. First, we tested this using direct co-culture. HUVECs plated on tissue culture plastic formed a typical monolayer (**Fig 3a**). When plated onto confluent CAF monolayers derived from female BC, they formed slender, longer tube-like structures reminiscent of vessels (**Fig 3b**). However, those plated on CAFs derived from male BC formed stubbier tube-like structures (**Fig 3c**) that, when quantified, were significantly shorter and with many branchpoints (**Fig 3d, e, respectively**).

**Figure 3:**
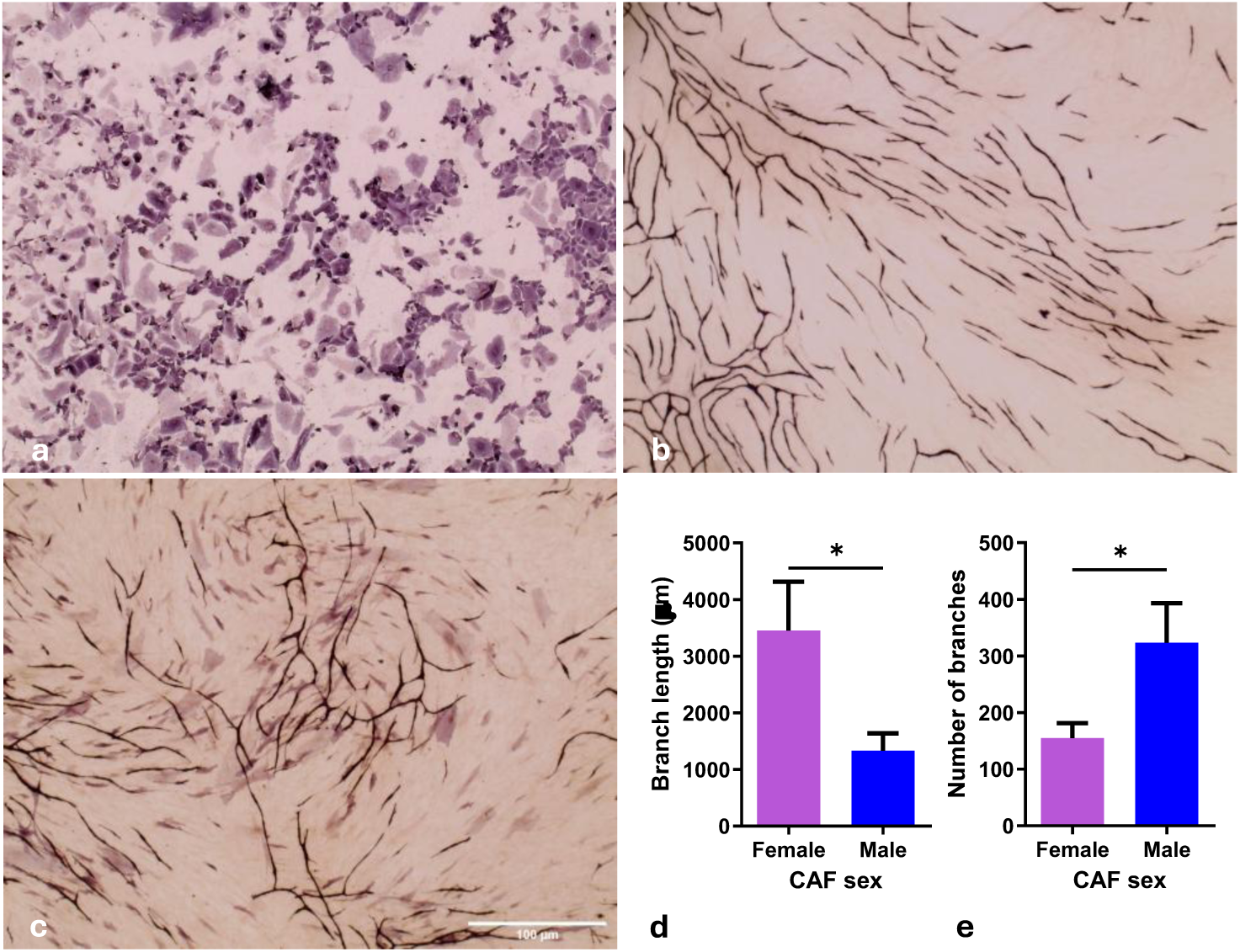
Induction of endothelial tube-like structures is dependent on CAF sex. HUVEC cultured alone formed a typical monolayer (a). Immunohistochemical analysis of CD31-stained cultures showed that HUVEC plated onto confluent female CAFs resulted in slender, longer vessels with little branching (b) whilst those on male CAFs formed fewer stubbier vessels with many branchpoints (c), quantified in d, e, respectively. * P <0.05. Scale bar = 100µ. Interestingly, a cell viability assay showed that HUVEC growth was not well supported by female CAF–CM, in contrast to CM from male CAFs (**Fig 4a**). However, in contrast, a wound closure assay showed that HUVECs closed the wound more quickly when exposed to CM derived from female BC-derived CAFs (**Fig 4b).** The presence of either male or female CAF CM had no significant impact on HUVEC motility speed (**Fig 4c**).

**Figure 4:**
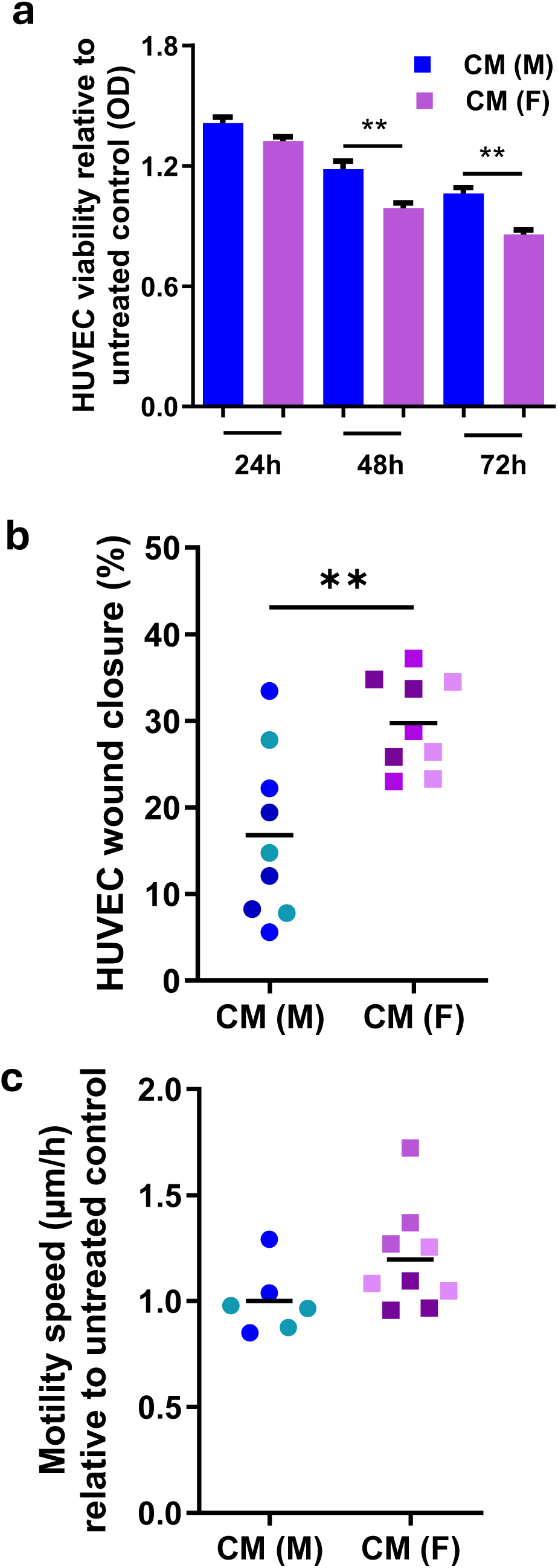
Effect of conditioned media (CM) generated from male and female breast cancer CAFs on HUVEC viability and wound closing ability. Cell viability was reduced in response to CM from female compared to male CAFs (a), while wound closure was accelerated female CAF CM (b). HUVEC motility was not affected by CAF CM from either sex. Values of CAFs from individual patient samples are indicated as coloured dots. ** P <0.001; * P<0.02.

These findings were extended to a microfluidic model to study vascular formation (**Fig 5a**). Microvascular networks were observed following HUVECs cultured alone (**Fig 5b**), or in the presence of female (**Fig 5c**) or male CAFs (**Fig 5d**). While some donor-dependent heterogeneity was observed, co-culture with female BC-derived CAFs promoted formation of enlarged channel-like structures and dense cellular aggregates (**Fig 5c).** In co-culture with male CAFs, HUVECs favoured interconnected capillary-like networks (**Fig 5d**). To explore how male or female CAFs influence the formation of vascular networks in the presence of cancer cells, MCF-7 cells were added to the microfluidic model. When MCF-7 cells were incorporated, the presence of female BC-derived CAFs supported well-structured vascular-like networks whereas male BC-derived CAFs showed less organised networks (**Fig 5e, f, respectively**). Together, these observations indicate the interaction between CAFs, endothelial cells and cancer cells may also be sex-dependent, paralleling observations from the CDM adhesion assays.

**Figure 5:**
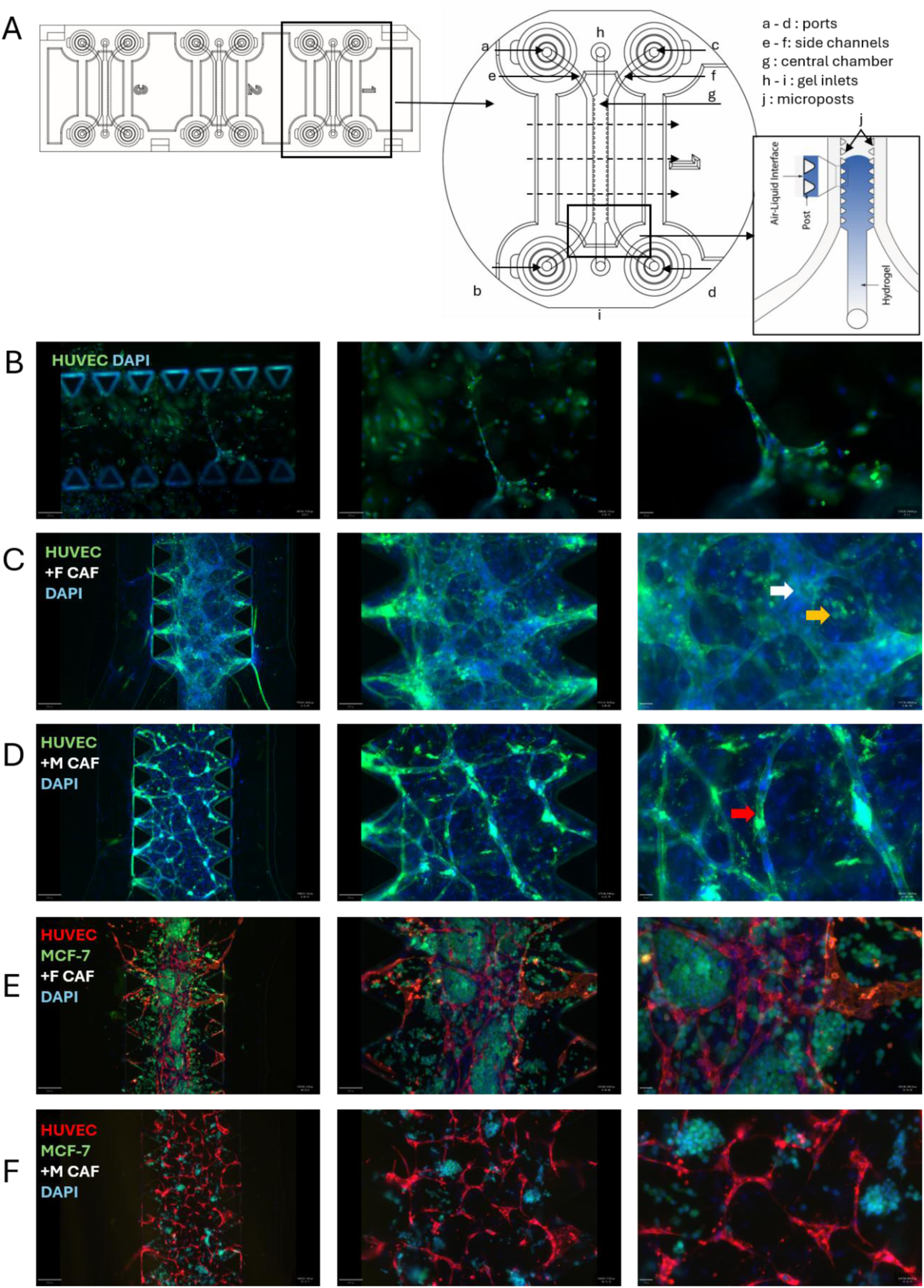
Microfluidic model of vasculature formation. Schematic diagram of the microfluidic device (a). Multiple configurations were tested to optimise fully animal-free culture conditions. Representative images show microvascular networks formed by HUVECs alone (b), HUVECs with female (c) and male BC-derived CAFs (d) after 14 days in culture. Female CAFs promoted the formation of enlarged hollow channels (white arrows) and dense cellular aggregates (yellow arrows), whereas male CAFs favoured interconnected capillary-like networks (red arrows). Incorporating MCF-7 cells supported well-structured vascular network in the presence of CAFs derived from female (e) but not male BC (f). Scale bars (e-f) = 750 µm, 300 µm, 150 µm (left to right.).

## Discussion

CAFs, the most abundant cell type within the TME, were regarded, historically as passive cells that provided purely structural support for tumours. Nowadays their wider roles in promoting cancer progression, therapy resistance and even immune evasion are recognised and they are themselves being mooted as possible targets for therapeutic intervention [42]. Even within BC subtypes, principal component analysis has revealed distinct gene expression profiles between luminal and HER2 subtypes with CAFs generated from the latter significantly enhancing the migration of T47D cells compared to the former [43]. CAF subtypes have also been identified in BC [26, 27]. Here, we have taken this a step further by demonstrating that CAFs derived from phenotypically matched male and female BC display distinct gene expression profiles when stratified by sex, and that this translated into biological function surrounding adhesion and vascularity. While sex-related biological outcomes have been identified in the cardiovascular field e.g. sex-dependent predisposition to atherosclerosis [44, 45] or as a direct response of endothelial cells [46] or fibroblasts [47] to external stimuli, importantly this study has identified that a specific cell type, the CAF, is a direct mediator of these differences in BC. Sexual dimorphism has been demonstrated in primary dermal fibroblasts isolated from non-sun-exposed skin of healthy young (<35 years) and aged (>55 years) male and female donors, supporting our findings in BC. Notably, aged male fibroblasts selectively promoted an invasive, therapy-resistant melanoma phenotype and enhanced metastatic progression in aged male mice [48]. Our data provides new evidence of sex-specific differences in BC CAFs, supporting recognised sex disparities in cancer epidemiology [49] and in the contribution of sex-biased transcriptional programs to cancer incidence [50].

Spatial analysis has allowed distribution patterns of CAFs to be generated, augmenting CAF phenotyping [27]. Different gene profiles are indicative that spatial architecture might shape CAF phenotypes. Our CAFs were generated from surplus breast tissue provided by pathology labs, meaning that defining spatial location was not possible. However, it is a reasonable assumption that bulk RNA-seq gene expression in these CAFs would reflect those identified from scRNA-seq data. Interestingly, when stratified for reported CAF subtypes, genes associated with iCAF and rCAF subtypes tended to be more closely associated with male CAFs, suggesting diverse origins from different cell precursors. While sex was not specifically mentioned, mining transcriptomic data from BC and adjacent normal tissues deposited in The Cancer Genome Atlas identified the iCAF phenotype as a poor prognostic marker [51].

Gene ontology analysis of RNA-seq data identified that the top enriched terms identified in male BC-derived CAFs were primarily associated with extracellular matrix organisation/adhesion and vascular development. This is consistent with another RNA-seq study which also compared CAFs from male and female BC and identified upregulation of genes involved in invasive and migratory processes [52]. The study established a nine-gene signature: *ASPN*, *COL4A1*, *COL4A2*, *COL4A5*, *COL5A3*, *COMP*, *EFEMP1*, *EMILIN2*, *FMOD*, *FN1*, *LAMA1* and *VTN* as drivers of the male phenotype [52]. While not tested in male BC, the nine-gene signature had strong prognostic value in prostate cancer. However, of these nine genes, only *COL5A3* and *COMP* were significantly upregulated in male BC-derived CAFs in our data. Other than stating that the nine-gene signature was generated from invasive mammary ductal carcinoma, detailed histopathological characterisation of the tissues that the CAFs were generated from was absent [52] hence the non-overlapping gene sets may have resulted from CAF-derivation from BCs of different histopathological type.

The physical appearance and phenotypic properties of CDMs revealed sex-specific differences in structure and thickness. In contrast to cell-free matrices derived from female 377HDFs and CDMs from female BC-derived CAFs, those from male BC-derived CAFs displayed a densely packed organisation comprising aligned fibres and was around one third thicker. This was supported by coherence analysis which revealed a linear alignment of collagen fibres in the male BC-derived CDMs. An aligned structure is consistent with facilitating the directional migration, invasion, and survival of cancer cells through so-called migration highways [53]. This phenotype has been observed in CAFs co-cultured with both premalignant and malignant mammary epithelial cells, but not in co-culture with benign fibroblasts from reduction mammoplasty [54]. Another study showed that cell-free matrices generated from normal fibroblasts and CAFs derived from the mammary tumour and healthy fat pad of FVB/n MMTV-PyMT mouse line, respectively were compositionally structurally distinct, with the latter stimulating cancer cell proliferation [55]. It is intriguing that male BC-derived CAFs displayed such a diverse CDM phenotype from both female CAFs and dermal fibroblasts in monoculture and points towards sex as a key variable.

Sex-dependency was also observed in cell adhesion to CDMs. Adhesion of non-transformed female mammary epithelial cells (HB2) did not differ between plastic or CDMs derived from male or female BC or from normal dermal fibroblasts. However non-transformed male mammary epithelial cells (MMEC) showed the highest level of adhesion of any cells tested, with particularly strong adhesion to male BC-derived CDMs. Male BC-derived CDMs also promoted superior adhesion of HUVEC. While HUVEC are generally provided commercially from umbilical cords from mixed sex, our findings with non-transformed mammary epithelial cells support the concept that sex-specific phenomena may occur independently of the whole-organism context [56]. In the context of whole tissue, fractal analysis of picrosirius red stained male and female BC tissue sections supported our *in vitro* observations, showing a more complex structural organisation in the collagen-rich TME of male BC, further pointing to sex-specific BC phenotypes. In the dental field, analysis of picrosirius red stained labial mucosal tissue showed greater proportion of thicker collagen fibres in male tissue [57]. Similarly, in animal models, collagen fibril diameter was related to sex, being significantly thicker in males [58].

Visually striking differences in sprouting angiogenesis and vessel geometry in response to CAF sex were observed. CAFs from male BC formed shorter, branched vessels, reminiscent of a disorganised angiogenic phenotype, where short, highly branched sprouts are indicative of increased endothelial branching activity but impaired vessel elongation and maturation. In a related angiogenesis assay, male human pulmonary microvascular endothelial cells produced fewer, longer, sprouts compared with those from females [46]. Phenotypic and functional differences between endothelial cells from diverse vascular pathologies is recognised [59]. Functional differences in vasculogenesis, extracellular matrix remodelling and angiogenic sprouting were observed between clinically prognostic CAF subtypes derived from pancreatic cancer in co-culture with HUVEC in a 3D microfluidic device, although CAF biological sex was not reported [60].

Sex-specific effects were also observed in CM experiments. The divergent effects of CM from different CAF sex may suggest functional heterogeneity between CAF populations, with male CAFs preferentially supporting tumour cell viability while suppressing processes involved in wound closure. The lack of differences in intrinsic motility may suggest that changes in wound healing could reflect altered proliferative capacity, cell–cell adhesion, or directional migration rather than a direct effect on migratory machinery. Additionally, CM may not necessarily increase how much a cell moves, but rather how efficiently it moves, shifting behaviour from random motility to persistent, directed migration [61].

As well as heterogeneity in responses to male versus female CAFs, we also observed heterogeneity in responses to CAF populations generated from different patients of the same sex. The heterogeneity of CAF populations is well recognised, for example, characterisation of CAFs from 14 female luminal BC patients revealed similar heterogeneity [62], consistent with the recognised heterogeneity of CAF populations and the varying abundance of specific CAF subtypes between individuals [27, 63]. This may reflect varying abundance of specific CAF subtypes between individuals [27, 63]. Interestingly our RNA-seq data hinted towards the presence of sex-specific CAF subtypes. ScRNA-sq analysis would be required to confirm if this truly reflects sex or relative abundance.

Finally, it has recently been shown that fetal bovine serum (FBS), a key additive to cell culture medium that promotes cell growth and survival, appeared to show sex-specific effects on gene expression and phenotype in cultured human endothelial cells and primary human lung fibroblasts [64]. This included enhanced vasculogenesis and vascular endothelial growth factor-induced sprouting in female cells, which was not seen in cells cultured under identical conditions but in the presence of charcoal-stripped serum which depletes FBS of hormones and growth factors. Hence, sex hormone effects from FBS may influence cellular responses. While an intriguing finding, the complexity and heterogeneity of FBS is recognised, with its biochemical composition incompletely characterised and substantial heterogeneity reported across numerous studies [65]. If proven by others, this may add further complexity to *in vitro* studies focused on the influence of sex. Importantly, our microfluidics experiments were conducted in the absence of any animal derived products, including no FBS and using recombinant human growth factors, suggesting true sex-dependent influences.

We have previously reported that male BC should be regarded as a separate BC subtype [6]. Our new in *vitro* findings clearly demonstrate a highly complex relationship between CAF sex and tumour characteristics, indicating that the most effective therapeutic approaches might be different for male and female BC. This has important implications for future research. Recognising that cellular sex is not reported in most published studies [66], our work highlights the need to integrate biological sex as a variable into pre-clinical BC studies. Further elucidation has potential to reveal sex-specific therapeutic strategies.

## Supporting information

Supplementary data Liu

## Acknowledgments

We are grateful to following for providing funding to support this work: Yorkshire Cancer Research, University of Aberdeen Development Trust, NHS Grampian Charity, Friends of ANCHOR, Breast Cancer Now, Animal Free Research UK and the Cyril & Margaret Gates Charitable Trust. The Breast Cancer Now Biobank provided some of the male-BC derived CAFs used in this study. Patient samples were obtained with appropriate ethical approval and informed consent. We thank those who kindly donated their tissue samples for use in this study.

## Author Contributions Statement

VS conceived and designed the study. PL and RAE contributed to the study design. PL, FRS, and VS contributed to data analysis, interpretation, and writing of the manuscript. PL, FRS, ME, NE, MPH, CCG, GC, LFS and RAE contributed to data collection and interpretation. Co-authors gave critical input, and all authors read and approved the final submitted version of the paper.

## Data Availability Statement

RNA-sequencing data have been deposited in the NCBI Gene Expression Omnibus (GEO) under accession number GSE337746 (https://www.ncbi.nlm.nih.gov/geo/). All other data supporting the findings of this study are included in the article and/or Supporting Information. Additional information is available from the corresponding author upon reasonable request.

