## Supplementary data Liu for "Sexual Dimorphism of Cancer-Associated Fibroblasts Governs Matrix and Vascular Organisation in Breast Cancer"

### Supplementary Figures

**Supplementary Figure S1** Generation and characterisation of CAFs

**Supplementary Figure S2** Expression CAF-, vascular- and epithelial associated genes in male and female BC-derived CAFs

**Supplementary Figure S3** CAF stratification according to published subtypes

**Supplementary Figure S4** Volcano plot

**Supplementary Figure S5** GO data

### Supplementary Tables

**Supplementary Table 1** CAF use in specific experiments

**Supplementary Table 2** Cells used and their culture conditions

**Supplementary Figures**

**Supplementary Figure S1** Generation and characterisation of CAFs

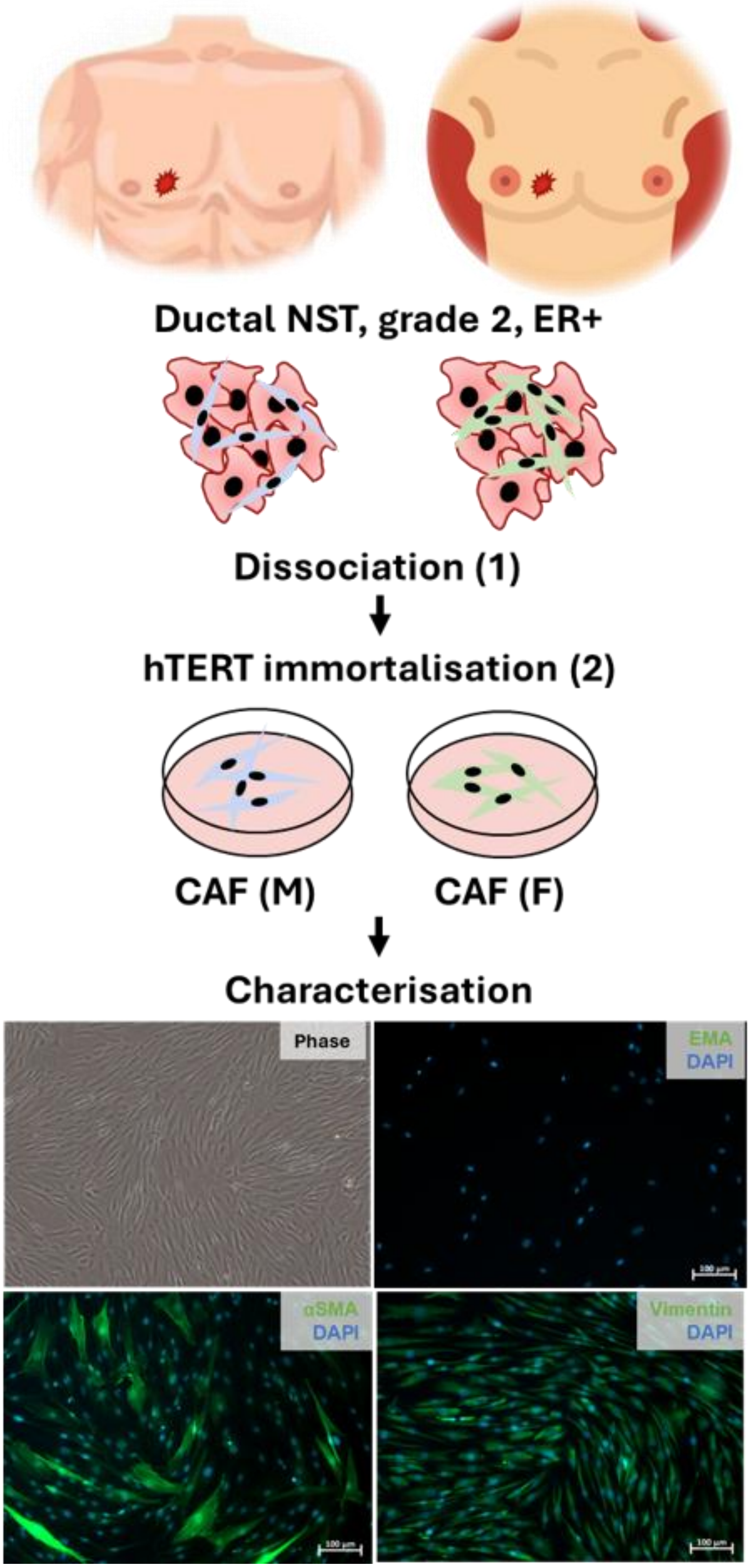


Pipeline for the generation of CAFs from male and female breast cancer. CAFs showed spindle shaped morphology and positivity for vimentin and α-SMA. Dissociation and hTERT immortalisation were achieved according to PMIDs: 9836473 and 16619045, respectively.

**Supplementary Figure S2** Expression CAF-, vascular- and epithelial associated genes in male and female BC-derived CAFs

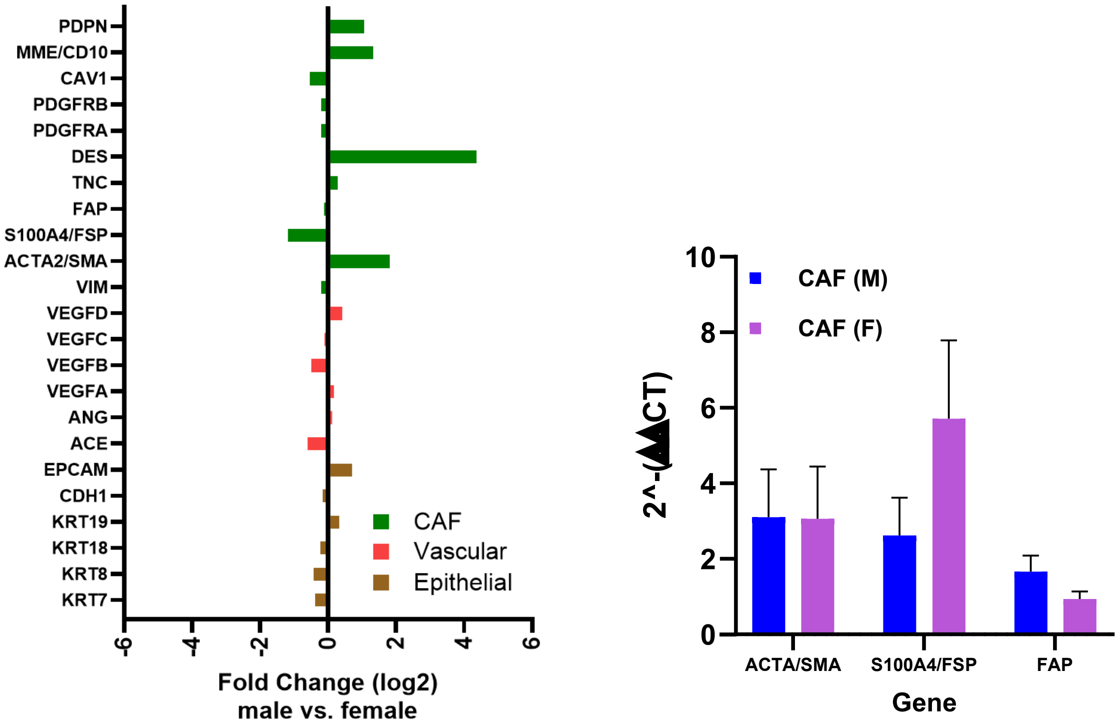


From RNA-seq data (left panel), the absolute $\log_2$ fold change between male and female BC-derived CAFs for recognised epithelial and vascular markers was around zero, indicating an absence of these cell types, with putative CAF markers expressed to varying levels between sexes, especially *DES*. Bars to the left represent genes overexpressed in CAFs derived from male breast cancer, those on the right are overexpressed in female breast cancer derived CAFs. qPCR validation of *ACTA/SMA*, *S100A4/FSP* and *FAP* are shown in the right panel.

**Supplementary Figure S3** CAF stratification according to published subtypes

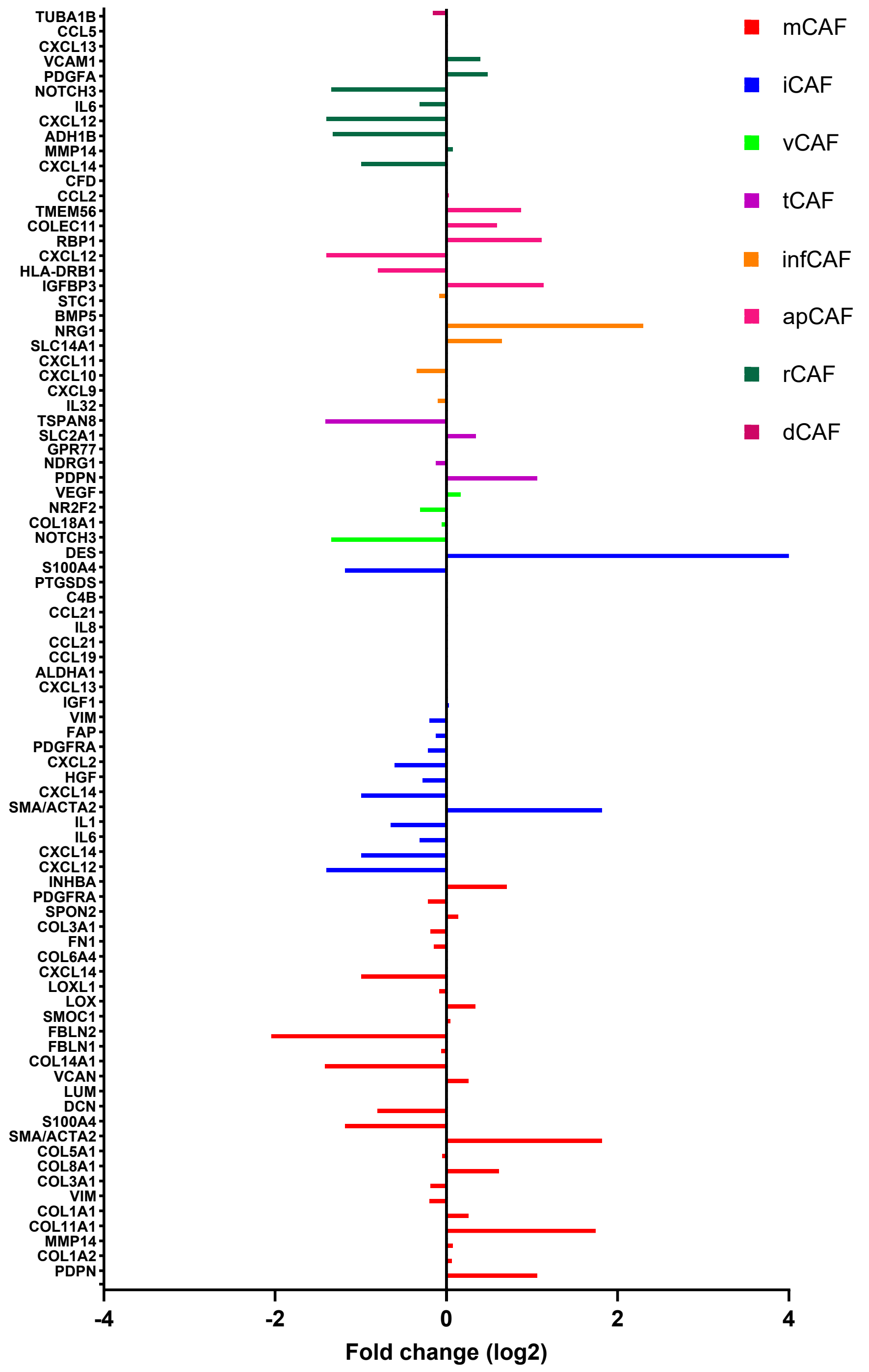


RNA-seq data generated from CAFs was categorised according to published marker genes that describe CAF types (PMID: 39255773). Genes to the left were overexpressed in CAFs derived from male breast cancer, those on the right were overexpressed in female breast cancer. mCAF, matrix CAF; iCAF, inflammatory CAF; vCAF, vascular CAF; tCAF, T-cell mediated CAF; infCAF, interferon CAF; apCAF, antigen-presenting CAF; rCAF, reticular-like CAF; dCAF, desmoplastic CAF.

**Supplementary Figure S4** Volcano plot

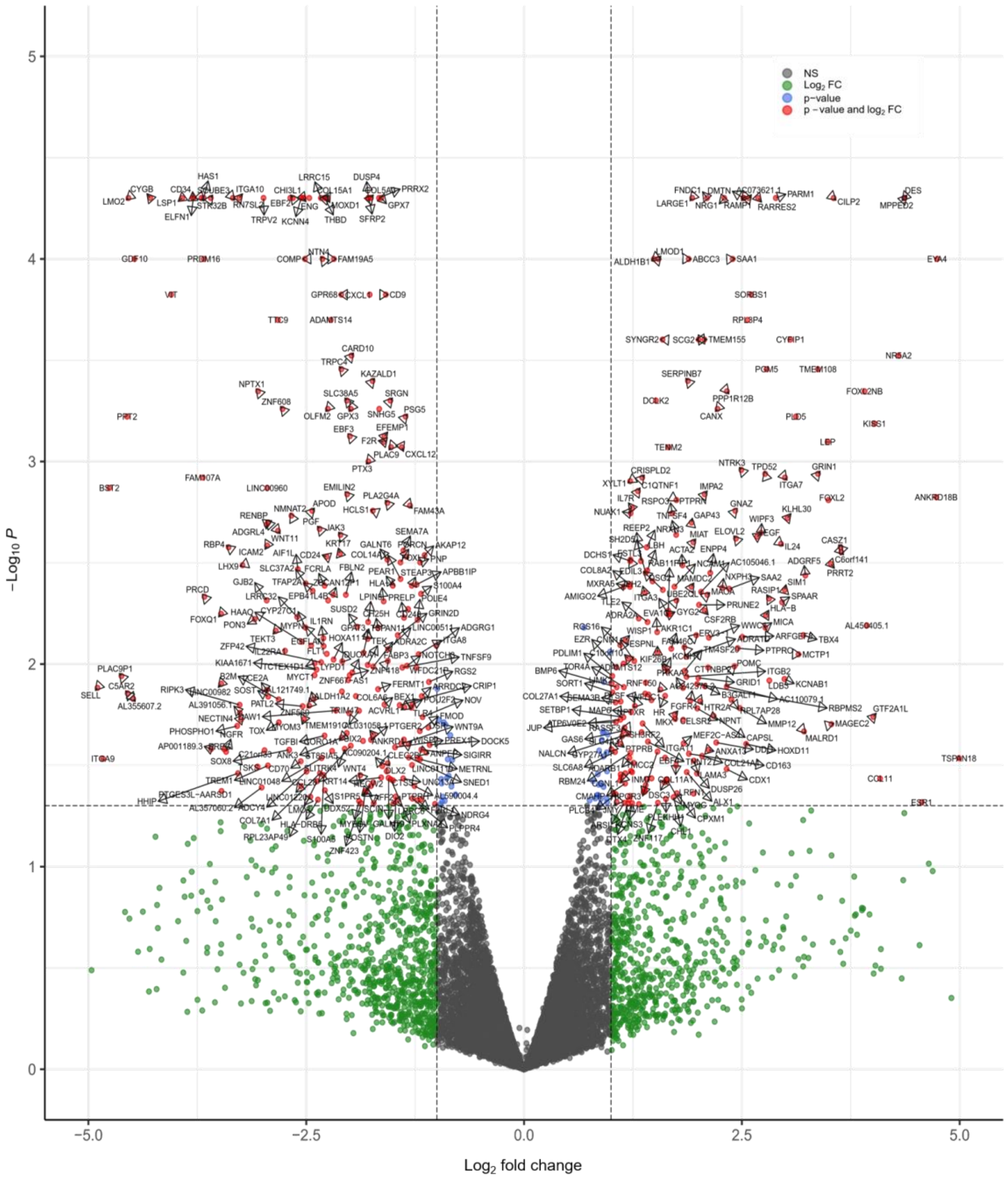


Volcano plot illustrating differential gene expression between CAFs derived from male and female BC. The x-axis shows the $\log_2$ fold change, and the y-axis shows the $-\log_{10}$(p-value). Genes significantly upregulated in male and female BC-derived CAFs are positioned towards the upper left and right, respectively. Significance was determined using adjusted p-values, with genes showing significance shown in red.

**Supplementary Figure S5** GO enrichment

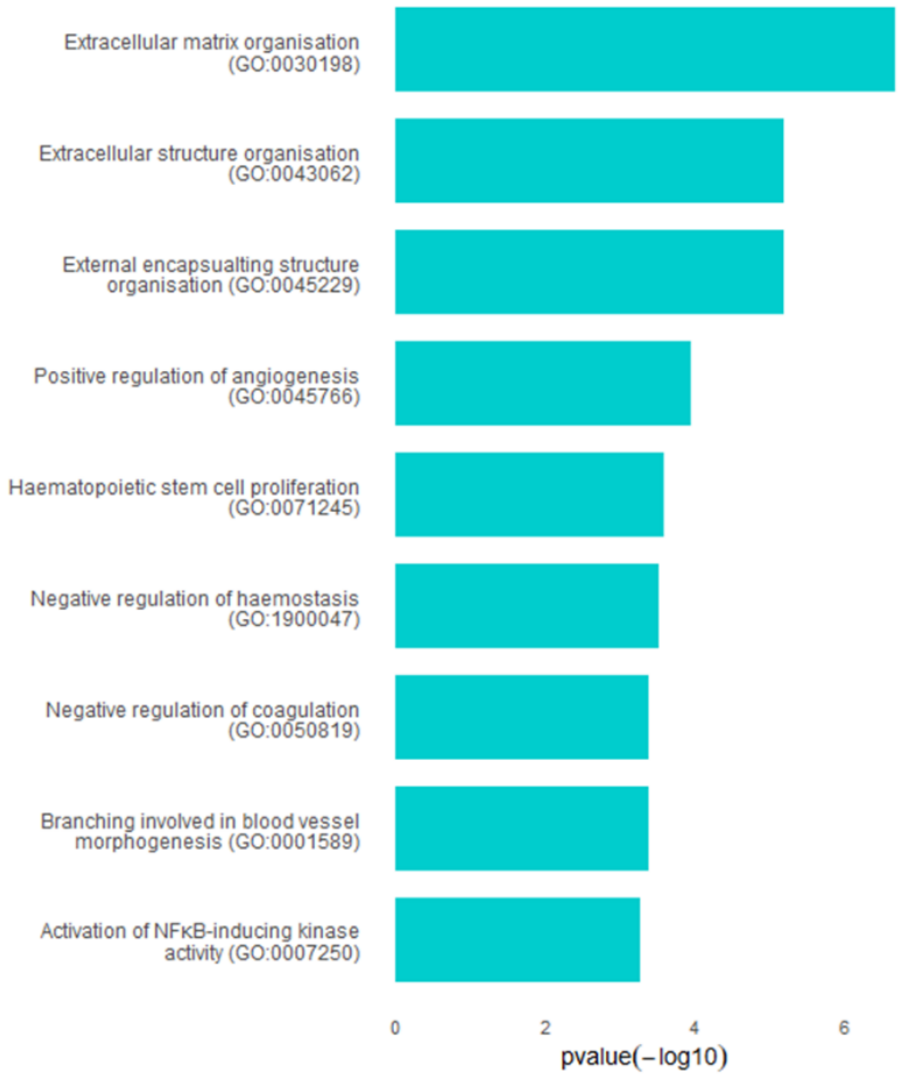


GO enrichment of DEGs in male BC-derived CAFs associated with matrix and vasculature identified using Enrichr (b).

**STable 1 CAF use in specific experiments**

| Case ID | Sex | RNA-seq | qPCR validation | Wound closure | Vessel formation | CM generation | Motility/migration | CDM generation | Adhesion assay | Tracking |
|---|---|---|---|---|---|---|---|---|---|---|
| LS10-210 | M | ✓ | ✓ | ✓ | ✓ | ✓ | ✓ | ✓ | ✓ | ✓ |
| LS15-02706 | M | ✓ | ✓ | ✓ | ✓ | ✓ | ✓ | ✓ | ✓ | ✓ |
| LS10-157* | M | ✓ | ✓ | ✓ | ✓ | | | | | |
| 2916T*# | M | | ✓ | | | | | | | |
| 1433T | M | | | | | | | | | |
| 1829T | M | | ✓ | | | | | ✓ | ✓ | ✓ |
| 1945T | M | | ✓ | | | ✓ | | ✓ | ✓ | ✓ |
| LS10-140 | F | ✓ | | | | | | | | |
| LS11-045 | F | | ✓ | ✓ | ✓ | ✓ | ✓ | ✓ | ✓ | |
| LS11-047 | F | | ✓ | ✓ | ✓ | ✓ | ✓ | ✓ | ✓ | ✓ |
| LS12-039 | F | ✓ | ✓ | | | | | ✓ | ✓ | |
| LS12-041 | F | ✓ | ✓ | ✓ | ✓ | ✓ | ✓ | ✓ | ✓ | ✓ |
| LS12-329 | F | | ✓ | ✓ | | | | ✓ | ✓ | ✓ |

CAFs were generated from age-matched ER-positive grade 2, lymph node positive ductal carcinoma of no specific type. CM, conditioned medium; CDM, CAF-derived matrix. *Stocks depleted, #non-immortalised.

**STable 2 Cells used and their culture conditions**

| Cell type | Description (source) | Culture conditions |
|---|---|---|
| CAFs | Human BC-derived carcinoma-associated fibroblasts (details in STable1) | DMEM high glucose with Glutamax™ (Gibco, UK), supplemented with with GlutaMAX™ and 10% (v/v) FBS (Bio-Sera, France) |
| HDF377 | Normal human dermal fibroblasts (Caltag Medsystems, UK) | DMEM high glucose with Glutamax™ (Gibco, UK), supplemented with 10% (v/v) FBS (Bio-Sera, France) |
| MCF-7 | Breast cancer cell line (ATCC/EACCC) | RPMI (Sigma, UK) supplemented with 5% (v/v) FBS (Bio-Sera, France) |
| HUVEC | Endothelial cells (cat#C12203 / C12253; Lot# 405Z020; Promocell, Germany) | Complete endothelial cell growth medium 2 (Promocell, Germany) |
| HB2 | Immortalised, non-transformed mammary epithelial cells (gift from Joyce Taylor-Papadimitriou, King's College London) | DMEM, high glucose with GlutaMAX™, supplemented with 10% FBS, 50 μg/mL hydrocortisone solution and 10 μg/mL human insulin |
| MEC | Male primary mammary epithelial cells (Lifeline Cell Technology, USA) | MammaryLife complete medium (cat#LL-0061) supplemented with recombinant human insulin (5 μg/mL), 6 mM L-glutamine, 1 μM Epinephrine, 5 μg/mL Apo-Transferrin, recombinant human TGF (0.5 ng/mL), 0.4% (v/v) Extract P™, Hydrocortisone Hemisuccinate 100 ng/mL (Lifeline Cell Technology, USA) |
